# Impaired cognitive performance under psycho-social stress in cannabis-dependent males is mediated by attenuated precuneus activity

**DOI:** 10.1101/429951

**Authors:** Weihua Zhao, Kaeli Zimmermann, Xinqi Zhou, Feng Zhou, Meina Fu, Christian Dernbach, Dirk Scheele, Bernd Weber, Monika Eckstein, René Hurlemann, Keith M Kendrick, Benjamin Becker

## Abstract

**Background:** Deficient regulation of stress plays an important role in the escalation of substance use, addiction and relapse. Accumulating evidence suggests dysregulations in cognitive and reward-related processes and the underlying neural circuitry in cannabis dependence. However, despite the important regulatory role of the endocannabinoid system in the stress response, associations between chronic cannabis use and altered stress processing on the neural level have not been systematically examined.

**Methods:** Against this background, the present functional magnetic resonance imaging (fMRI)study examined psycho-social stress processing in cannabis-dependent males (n = 28) and matched controls (n = 23) using an established stress-induction paradigm (Montreal Imaging Stress Task) that combines computerized (adaptive) mental arithmetic challenges with social evaluative threat.

**Results:** During psycho-social stress exposure, but not the no-stress condition, cannabis users demonstrated impaired performance relative to controls. In contrast, levels of experienced stress and cardiovascular stress responsivity did not differ from controls. Functional MRI data revealed that stress-induced performance deteriorations in cannabis users were accompanied by decreased precuneus activity and increased connectivity of this region with the dorsal medial prefrontal cortex.

**Limitations:** Only male cannabis-dependent users were examined, the generalizability in female users remains to be determined.

**Conclusion:** Together, the present findings provide first evidence for exaggerated stress-induced cognitive performance deteriorations in cannabis users. The neural data suggest that deficient stress-related dynamics of the precuneus may mediate the deterioration of performance on the behavioral level.

## Introduction

Drug addiction has been conceptualized as a chronic and relapsing brain disorder. On the symptomatic level, the condition is characterized by escalating drug intake, progressive loss of behavioral control, withdrawal and strong craving in response to drug-cues or stressors.^1^ Current neurobiological perspectives propose that the transition from occasional to addictive drug use is accompanied by progressive mal-adaptations in neural circuits engaged in reward processing, associative learning, executive control, and stress reactivity.^2^

Cannabis is the most widely used illicit drug, with 3.8% of the world’s population consuming cannabis on a regular basis.^3^ Cannabis use-associated alterations in the domains of reward processing and cognition have been extensively studied^4,5^ and there is growing evidence from functional imaging studies suggesting neuroplastic adaptations in neural systems subserving these functions.^6,7^ In the cognitive domain, selective impairments in attention, working and associative memory have been reported most consistently.^4,5,8,9^

Long-term stress has detrimental effects on mental health^10^ and the acute stress response is an adaptive mechanism to environmental demands that are perceived as potentially threatening. Deficient regulation of stress is considered a hallmark of addiction,^11,12^ and represents a risk-factor for both, the escalation of drug use^13–15^ and relapse.^12,16^ Impairments in the neural circuits that mediate the acute adaptive response may contribute to reduced access to adaptive coping, such that cognitive functions, including sustained attention and regulatory control, become compromised.^16,17^

A growing number of functional imaging studies reported abnormal emotional reactivity and emotion regulation in cannabis users, including abnormal neural reactivity to affective stimuli^18,19^ and deficient amygdala downregulation during cognitive reappraisal.^20^ However, despite accumulating evidence for altered emotional reactivity and cognitive emotion regulation, it remains unknown whether deficient stress regulation may contribute to cannabis dependence. Support for an association between deficient stress regulation and cannabis dependence comes from large-scale surveys reporting that regulation of negative affect represents a primary motivation for cannabis use^21^ and that this coping-oriented motivation increases the risk to develop patterns of dependent use.^22^

To determine the integrity of the behavioral and neural stress responses in cannabis dependence, the present study administered an established psychosocial stress induction fMRI paradigm (Montreal Imaging Stress Task, MIST) to cannabis-dependent males and matched non-using controls. During the MIST, subjects are required to perform an adaptive arithmetic task combined with negative performance feedback and critical social evaluation. To additionally examine the effects of stress on the urge to use cannabis, craving was assessed before and after psychosocial stress induction. Based on previous studies, we expected that cannabis-dependent participants would exhibit an impaired stress regulation capacity in the context of altered neural activity in circuits mediating psycho-social stress processing.

## Methods

### Participants

*N* = 34 cannabis dependent males and *n* = 28 non-using healthy controls were recruited in cooperation with local drug counseling centers and by additional advertisements. All cannabis users fulfilled the DSM-IV criteria for cannabis dependence (Mini-International Neuropsychiatric Interview, MINI).^23^ To reduce non-cannabis use associated variance the study enrolled only male participants. The decision was based on previous studies reporting sex-differences during psycho-social stress induction,^24^ stress induced drug-craving^25^ and menstrual cycle effects on emotion regulation.^26^ A similar approach has been employed in previous studies on stress reactivity^27^ and in previous studies examining emotional processing in cannabis users.^19,20,28,29^

Exclusion criteria included (1) age < 18 or > 40 years, (2) left-handedness, (3) history or current DSM-IV axis I disorders (based on MINI, exception: cannabis abuse or dependence), (4) Beck Depression Inventory (BDI-II) score > 20^30^, (5) current or history of a medical disorder including endocrinological abnormalities (6) current or regular use of medication, (7) usage of other illicit substances > 75 lifetime occasions or during the 28 days prior to the experiment, (8) positive urine screen for cocaine (300 ng/ml), methamphetamine (500 ng/ml), amphetamine (500 ng/m), methadone (300 ng/ml) or opiate (300 ng/ml) (Drug-Screen-Multi 7TF, von minden GmbH, Moers, Germany), (9) breath alcohol >0.00% (analyzed using TM-7500, Trendmedic, Penzberg, Germany). For controls, additional exclusion criteria were applied: (1) cumulative lifetime use of cannabis > 15 g (*M* = 1.29, *SD* = 1.02), (2) use of any other illicit substance >10 lifetime occasions. To control for confounding sub-acute effects of cannabis, all users were required to remain abstinent from cannabis 24h prior to the fMRI experiment. To increase adherence with the abstinence period participants were informed that a urinary drug test for cannabis use would be performed on the day of the experiment.^31,32^

Screening procedures and fMRI assessment were scheduled on separate study days. Following an initial brief telephone screening, potential eligible participants were invited for an in-depth screening that included assessment of study participation criteria (i.e. diagnostic interview using the MINI, drug use interviews). Eligible subjects were scheduled for the fMRI assessment and informed about the required abstinence time. The fMRI assessment was preceded by the assessment of potential confounders, including drug screenings (i.e. urinary drug test, breath alcohol test) as well as assessment of emotional state and baseline cognitive performance indices, specifically anxiety (State Trait Anxiety Inventory, STAI),^33^ mood (Positive and Negative Affect Schedule, PANAS)^34^ and attention (d2 test of attention).^35^ Given the high prevalence of tobacco co-use in cannabis dependence^36^ the experimental groups were matched in terms of nicotine use. However, both, acute nicotine administration and abstinence-induced nicotine craving may impact stress processing and underlying neural mechanisms.^37^ Nicotine craving is reported to peak around 3-6 hours following the last cigarette and a recent study reported craving-associated neural activity changes after 4h of abstinence.^38^ As a trade-off, subjects were allowed to smoke as usual, however underwent a 1.5h supervised abstinence period before the start of the experimental paradigm.^20,29^ Following initial quality assessments 6 cannabis users and 5 controls were excluded (supplemental materials, Figure S1), the final dataset included *n* = 28 dependent cannabis users and *n* = 23 healthy controls.

All subjects provided written informed consent and the study procedures had full approval by the local ethics committee at the University of Bonn, Germany. All study procedures were in accordance with the latest revision of the Declaration of Helsinki.

### Experimental design

Psychosocial stress during fMRI acquisition was induced using the MIST.^39^ During the paradigm, subjects were asked to perform mental arithmetic tasks and were confronted with negative feedback about their performance indicating that they performed worse than the other study participants.^27^ Briefly, before the experiment subjects were instructed to perform the task with high accuracy and speed. The instructions emphasized that it would be very important that subjects match the average performance of the other participants and that the experimenters would monitor and evaluate the performance online via monitor. To further increase the psychosocial stress the experimenters criticized the participants’ “bad” performance via intercom and reminded them of the importance to perform similar to the other participants between the runs of the task.

The block-design fMRI paradigm consisted of six runs (each 6 minutes): three runs with negative feedback (stress condition) and three runs without feedback (no-stress control condition). The order of runs was fixed (a no-stress run always followed by a stress run). Each run incorporated four 60 s blocks that were preceded by a visual attention cue (5 s) and followed by a 20 s inter-block interval that served as low-level baseline (fixation cross). During the blocks, subjects were required to perform an arithmetic task and to select the correct answer using a rotary dial. The subjects received feedback (“correct” or “incorrect”) on whether their response was correct or incorrect. During the stress blocks, additional performance indicators were displayed (own performance and average performance of the other subjects). To further increase stress, a time limit was implemented that was indicated by a progressing bar moving from the left to right and “time out” was displayed, in case no response occurred during the given time. Unbeknownst to the subject, an algorithm was employed that adopted the response times to the performance of the subject to increase the failure rate. First, the average response time of the participant was determined in a pre-scan training session of 2 minutes without a time limit per arithmetic task and the time limit for the task during fMRI was set to 90% of the subject’s individual baseline response time. Furthermore, the time limit was decreased by 10% after three correct responses and likewise increased by 10% after three incorrect responses.^39^

After each run subjects rated their stress level on a scale from 1 (very low) to 8 (very high). To determine baseline and stress-induced cannabis craving, all subjects rated their level of cannabis craving (visual analog scale, VAS, 0-100) before and after the paradigm. To control for between-group differences in task engagement and self-perceived performance, subjects rated task enjoyment (1-9 points, 1 - very unpleasant, 9 - very pleasant) and their own performance (1-9 points, 1 - very negative, 9 - very positive) at the end of the experiment. As a physiological indicator of stress, blood pressure (systolic and diastolic) was repeatedly measured at four different time points (at rest 30 minutes after arrival - t1, immediately before the task - t2, immediately and 60 minutes after the task - t3 and t4). Blood pressure data for five subjects was lost due to technical failure, leading to a final sample size of *n* = 23 controls and *n* = 23 users for the corresponding analysis.

The experiment lasted approximately 40 minutes. Stimuli were presented via liquid crystal display video goggles (Nordic NeuroLab, Bergen, Norway).

### fMRI acquisition and processing

Data was acquired at 3Tesla and preprocessed using standard protocols in SPM12 (Wellcome Department of Imaging Neuroscience, UCL, London, UK). The first level design matrix employed a boxcar function to model the ‘stress’ and ‘no-stress’ condition and additionally included head motion parameters as nuisance regressors (details provided in supplemental materials).

### fMRI BOLD level analyses

On the second level, we first investigated the stress-network using a one sample t-test on the pooled data from cannabis users and controls (stress > no-stress). To determine altered neural stress processing in cannabis users, a two-sample t-test was conducted comparing stress-related activity between the groups (stress > no-stress). To increase the sensitivity of the analysis, the task-specific stress neural networks were initially defined with an independent data set from a previous study using the identical paradigm.^27^ Briefly, to independently determine the task-specific stress network, fMRI data from this independent sample of *n* = 31 healthy male participants (only non-treatment subjects from the previous study)^27^ was subjected to a one-sample t-test in SPM using the (stress > no-stress) contrast and thresholded at *p* < 0.05 using family-wise error correction (FWE). Results revealed that the paradigm significantly engaged the middle temporal gyrus, precuneus, (para-)hippocampal gyrus and inferior parietal lobule which were consequently considered as stress sensitive regions. Based on these results, bilateral anatomical masks of these four regions as provided in the Automatic Anatomical Labeling (AAL) atlas implemented in the WFU PickAtlas^40^ were combined into a single mask. This mask was subsequently used for small volume correction (SVC) employing an FWE-corrected *p* < 0.05. For further post hoc analyses parameter estimates were extracted from spheres with 6 mm radius centered at the maximum t-value coordinates of between-group differences using MarsBar.^41^

### fMRI functional connectivity analyses

To further explore whether neural activity alterations in cannabis users were associated with altered network-level communication, a generalized form of context-dependent psychophysiological interaction (gPPI) analysis was performed.^42^ To this end, the task-related functional connectivity of regions that exhibited significant between-group differences in the BOLD-level analysis were examined (precuneus, see Results – fMRI BOLD level). The first level gPPI models were modelled after deconvolution and included a psychological factor, physiological factor and the interaction between the two factors (PPI term). In addition to the experimental conditions head motion parameters were included as nuisance regressors. In line with the BOLD-level analysis, differences between cannabis users and controls were determined by means of two-sample t-tests in SPM using the contrast (stress > no-stress). Between-group differences in task-related functional connectivity were determined on the whole-brain level using *p* < 0.001 uncorrected (only clusters with minimum voxel size >10 reported). Given that between-group differences in task-related functional connectivity were examined using an uncorrected threshold, the corresponding findings are considered as exploratory.

## Results

### Potential confounders and drug use patterns

The groups were comparable with respect to several important confounders (*ps* > 0.10). Use of other prevalent illicit drugs was low in both groups, however, cannabis users reported more occasions of ecstasy use (*M* ± *SD*: 9.72 ± 2.19 vs. 2.33 ± 2.31). Details and cannabis use parameters provided in **Table 1**.

**Table 1.** Group characteristics and substance use (*n* = 51).

| Measurements | Control (*n* = 23) | User (*n* = 28) | *t* ($\chi^2$) | *p* |
| --- | --- | --- | --- | --- |
| | mean ± SD | mean ± SD | | |
| Age (years) | 24.57±3.55 | 25.54±5.11 | 0.77 | 0.45 |
| Education (years) | 16.20±2.28 | 15.95±3.68 | 0.30 | 0.77 |
| Number of smokers | 19 | 26 | 1.09* | 0.30 |
| Years of nicotine use | 7.21±2.94 | 9.65±5.75 | 1.68 | 0.10 |
| Cigarettes per day | 8.45±5.34 | 9.51±7.11 | 0.57 | 0.57 |
| Packyears | 3.04±2.44 | 5.56±6.33 | 1.63 | 0.11 |
| Onset of alcohol use | 16.57±1.85 | 15.98±3.80 | 0.66 | 0.51 |
| Days per week with alcohol use | 1.24±1.05 | 1.56±1.35 | 0.90 | 0.37 |
| Quantity (units) of alcohol per week | 6.31±5.67 | 10.14±9.01 | 1.70 | 0.10 |
| Fagerström test, Nicotine dependence | 1.00±1.31 | 1.86±2.22 | 1.63 | 0.11 |
| Age of first nicotine use | 15.71±1.74 | 14.96±2.49 | 1.22 | 0.23 |
| PANAS P | 21.17±5.51 | 20.00±6.43 | 0.69 | 0.49 |
| PANAS N | 1.96±3.14 | 1.29±2.75 | 0.81 | 0.42 |
| BDI II | 6.09±6.26 | 6.50±4.42 | 0.28 | 0.78 |
| STAI -State | 32.83±5.69 | 31.71±5.77 | 0.69 | 0.49 |
| STAI -Trait | 35.48±8.17 | 35.86±7.71 | 0.17 | 0.87 |
| Age at first use of cannabis | 16.92±2.57 | 15.68±2.82 | 1.51 | 0.14 |
| Average frequency<br>(last 12 months, days per month) | - | 23.61±7.27 | - | - |
| Lifetime quantity (g) | 1.29±1.02 | 2309±1655 | 7.38 | 0.00 |
|  |  |  |  | 1 |
| Participants with past ecstasy use | 3 | 18 | - | - |
| Occasions of ecstasy use | 2.33±2.31 | 9.72±2.19 | 2.88 | 0.01 |
| Participants with past cocaine use | 2 | 14 | - | - |
| Occasions of cocaine use | 5.50±4.50 | 5.93±2.73 | 0.08 | 0.94 |
| Participants with past amphetamine use | 4 | 18 | - | - |
| Occasions of amphetamine use | 3.25±4.50 | 12.17±15.39 | 1.13 | 0.27 |
| Participants with past hallucinogen use | - | 17 | - | - |
| Occasions of hallucinogen use | - |  | - | - |
| Participants with past opiate use | - | 3 | - | - |
| Occasions of opiate use | - |  | - | - |
| Participants with past solvents use | - | 7 | - | - |
| Occasions of solvents use | - |  | - | - |
| Post experiment |  |  |  |  |
| Task pleasantness | 6.09±1.93 | 6.11±1.99 | 0.04 | 0.97 |
| Evaluation of performance | 5.87±1.33 | 5.18±1.66 | 1.62 | 0.11 |

SD = standard deviation
\* $\chi^2$ test employed

### Craving, stress experience, performance and blood pressure

Examining cannabis craving using a mixed ANOVA with group (control vs. user) as a between subject factor and time (pre-vs. post-stress task) as within-subject factor revealed significant main effects of group [*F*_(1, 49)_ = 51.20, *p* < 0.001, partial *η^2^* = 0.51] and time [*F*_(1, 49)_ = 6.23, *p* = 0.02, partial *η^2^* = 0.11], and a significant interaction effect [*F*_(1, 49)_ = 4.23, *p* = 0.05, partial *η^2^*= 0.08]. Post-hoc analyses showed generally higher craving ratings in the cannabis group. Importantly, within the group of cannabis users craving strongly increased after stress exposure (cannabis user group, *p* = 0.001, Cohen’s *d* = 0.38, control group, *p* = 0.77, see **Figure 1A**).

**Figure 1.**
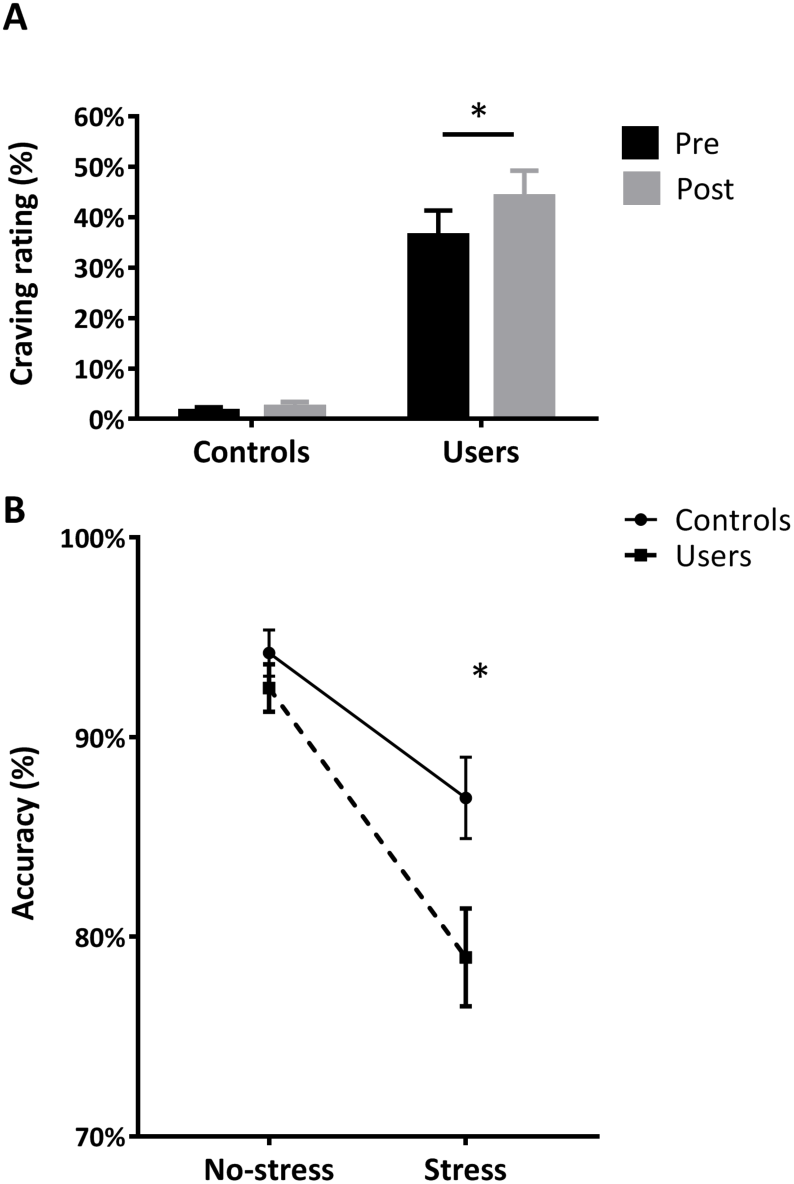
**(A)** Cannabis craving assessed before and after stress-induction, **(B)** performance accuracy during the no-stress and stress condition. Error bars reflect the SEM. \**p* < 0.05.

Stress experience was analyzed by means of a mixed ANOVA with the between subject factor group (control vs. user) and the within subject factor condition (stress vs. no-stress). The main effect of condition was significant [*F*_(1, 49)_ = 131.91, *p* < 0.001, partial *η*^2^ = 0.73] indicating successful stress-induction. However, there was no significant interaction effect indicating that both groups experienced comparable levels of subjective stress.

Examining accuracy (percent correct responses) using a concordant mixed ANOVA revealed a main effects of condition [*F*_(1, 49)_ = 60.00, *p* < 0.001, partial *η^2^* = 0.55] reflecting that both groups performed better during the no-stress condition. Furthermore, there were a main effect of group [*F*_(1, 49)_ = 4.54, *p* = 0.04, partial *η^2^* = 0.09] and a significant group x condition interaction effect [*F*_(1, 49)_ = 5.15, *p* = 0.03, partial *η^2^* = 0.10]. Bonferroni-corrected post-hoc tests revealed that the groups exhibited comparable performance during the no-stress condition (*p* = 0.31). However, under stress cannabis users performed significantly worse than controls (*p* = 0.02, Cohen’s *d* = 0.65, see **Figure 1B**).

Examination of blood pressure revealed a significant main effect of time (t1/ t2/ t3/ t4) for systolic [*F*_(3, 132)_ = 6.19, *p* = 0.001, partial *η^2^* = 0.12] and diastolic blood pressure [*F*_(3, 132)_ = 4.63, *p* = 0.005, partial *η^2^* = 0.10]. Bonferroni-corrected pairwise comparisons illustrated that systolic blood pressure after the task was higher compared to 30 minutes after arrival (*p* = 0.03) and immediately before the task (*p* = 0.001) reflecting successful stress induction. Diastolic blood pressure was higher at t3 compared to t2 (*p* = 0.006). In line with the lack of between-group differences in self-reported stress experience, both groups displayed comparable cardio-vascular stress reactivity.

### fMRI – BOLD level

In line with previous studies^27^, the paradigm induced widespread activity in psycho-social stress networks encompassing middle frontal regions, precuneus and posterior cingulate cortex (**Table 2**, **Figure S2**). Examining neural differences between the cannabis users and controls in the task-specific stress network revealed significantly decreased stress-reactivity in the cannabis users as compared to controls in a cluster located in the right precuneus (3, - 70, 50, *k* = 32, FWE *p* < 0.05). Post-hoc analyses of extracted parameter estimates from this region further revealed that the differences during stress vs. no-stress in precuneus were smaller in cannabis users relative to controls [*t* _(49)_ = 2.90, *p* = 0.006, Cohen’s *d* = 0.8, **Figure 2C**] and cannabis users showed an attenuated increase during stress relative to the no-stress condition (*t* _(27)_ = 2.47, *p* = 0.023, Cohen’s *d* = 0.32, corresponding to a small effect size) as compared to controls (*t* _(23)_ = 5.71, *p* < 0.001, Cohen’s *d* = 0.83, corresponding to a large effect size) (**Figure 2D**). This was also reflected by marginally significant lower precuneus activity during stress in the cannabis users as compared to controls (*t* _(49)_ = 1.74, *p* = 0.088) (**Figure 2D**). An exploratory whole brain analysis did not reveal significant between group differences on the whole brain level after FWE-correction.

**Figure 2.**
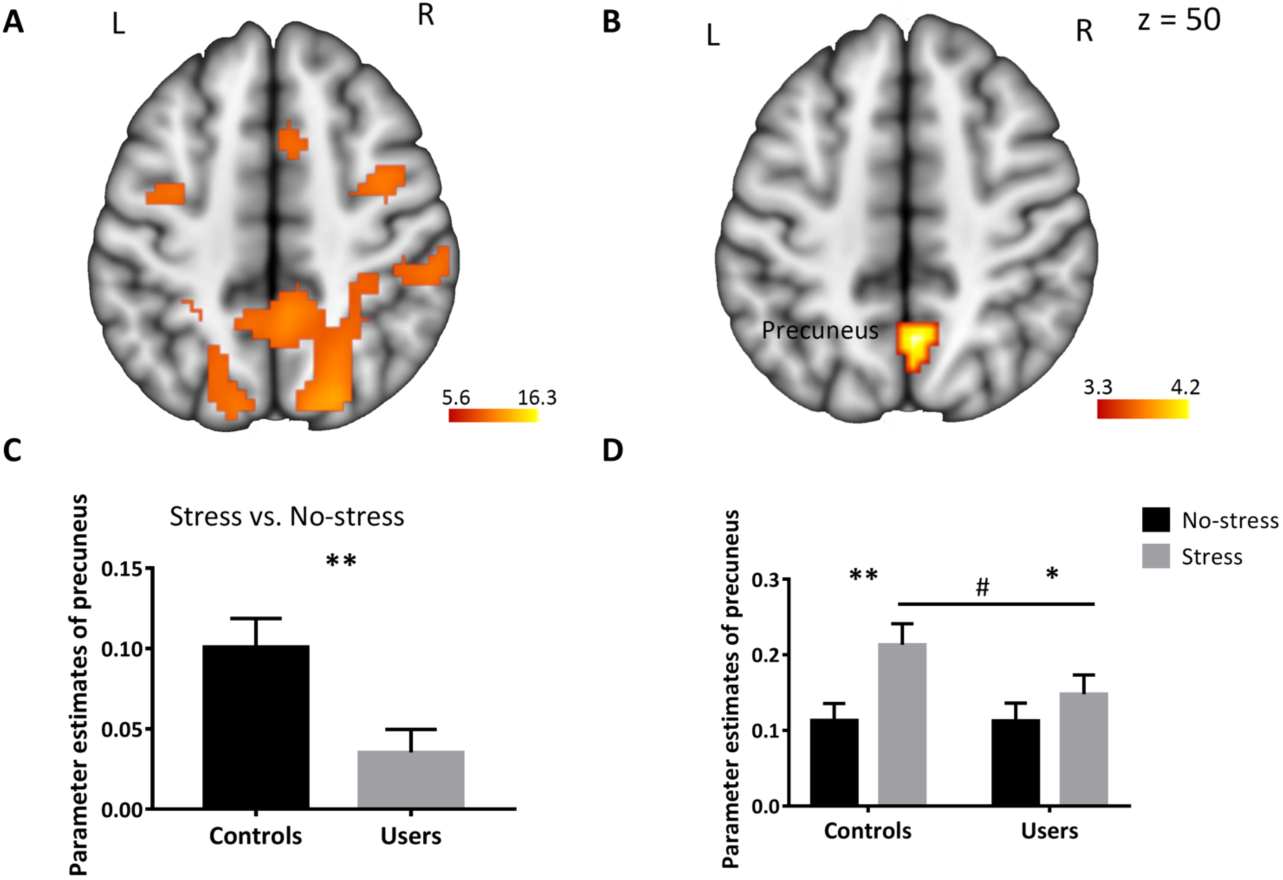
Stress-related network in the entire sample (**A**) and differences between cannabis users and controls (**B**) showing that cannabis users exhibited decreased precuneus activity during psycho-social stress. Extracted parameter estimates from the precuneus for the contrasts (stress > no-stress) **(C)** and (no-stress vs baseline, stress vs baseline) **(D)** further revealed that the effect was driven by lower activity during the stress condition. Image (A) is thresholded at *p* < 0.05, family-wise error corrected (FWE). Image (B) is thresholded at *p*_FWE_ < 0.05, small volume correction. Error bars reflect the SEM. \**p* < 0.05. \*\**p* < 0.01. ^#^ *p* = 0.088. Laterality [left (L), right (R)].

**Table 2.** Stress-related activity increases in the entire sample (Stress > No-stress)

| Hemisphere | Region | Cluster size | Peak t | MNI coordinates |  |  |
| --- | --- | --- | --- | --- | --- | --- |
|  |  |  |  | x | y | z |
| Stress > No-stress |  |  |  |  |  |  |
| R | Precuneus/ Middle Occipital Gyrus | 6864 | 16.30 | 9 | -76 | -10 |
|  |  |  | 16.04 | 45 | -64 | 8 |
|  |  |  | 15.69 | 18 | -94 | 14 |
| R | Insula/ Middle Frontal Gyrus | 400 | 10.48 | 33 | 20 | 5 |
|  |  |  | 7.73 | 33 | 44 | 20 |
|  |  |  | 7.61 | 33 | 41 | 32 |
| R | Middle Frontal Gyrus/ Medial Frontal Gyrus/ Cingulate Gyrus | 603 | 9.72 | 39 | -1 | 53 |
|  |  |  | 9.14 | 45 | 8 | 32 |
|  |  |  | 8.58 | 21 | -1 | 59 |
| L | Middle Frontal Gyrus | 69 | 7.50 | -39 | -10 | 50 |

Stress-related activity increases in the entire sample (Stress > No- stress). The table gives anatomical regions, cluster size, peak t value, their laterality [left (L), right (R)], and MNI coordinates (x, y, z). Laterality [left (L), right (R)].

### Associations between neural activity and cannabis use parameters

In the cannabis dependent group, no significant associations between precuneus activity and cannabis use parameters (age of onset, cumulative lifetime use, frequency of use) were observed *p*s > 0.22.

### Functional connectivity

An exploratory whole-brain analysis comparing stress-related connectivity of the precuneus (seed region) between cannabis users and controls revealed relatively increased connectivity of the precuneus with the dorsal medial prefrontal cortex (dmPFC; 12, 26, 56, *k* = 34, *p* < 0.001, uncorrected) in the cannabis users (**Figure 3A**) and independent *t*-test suggested the precuneus-dmPFC coupling was stronger in cannabis users compared to controls [*t* _(49)_ = 3.62, *p* = 0.001, Cohen’s *d* = 1.02, **Figure 3B**]

**Figure 3.**
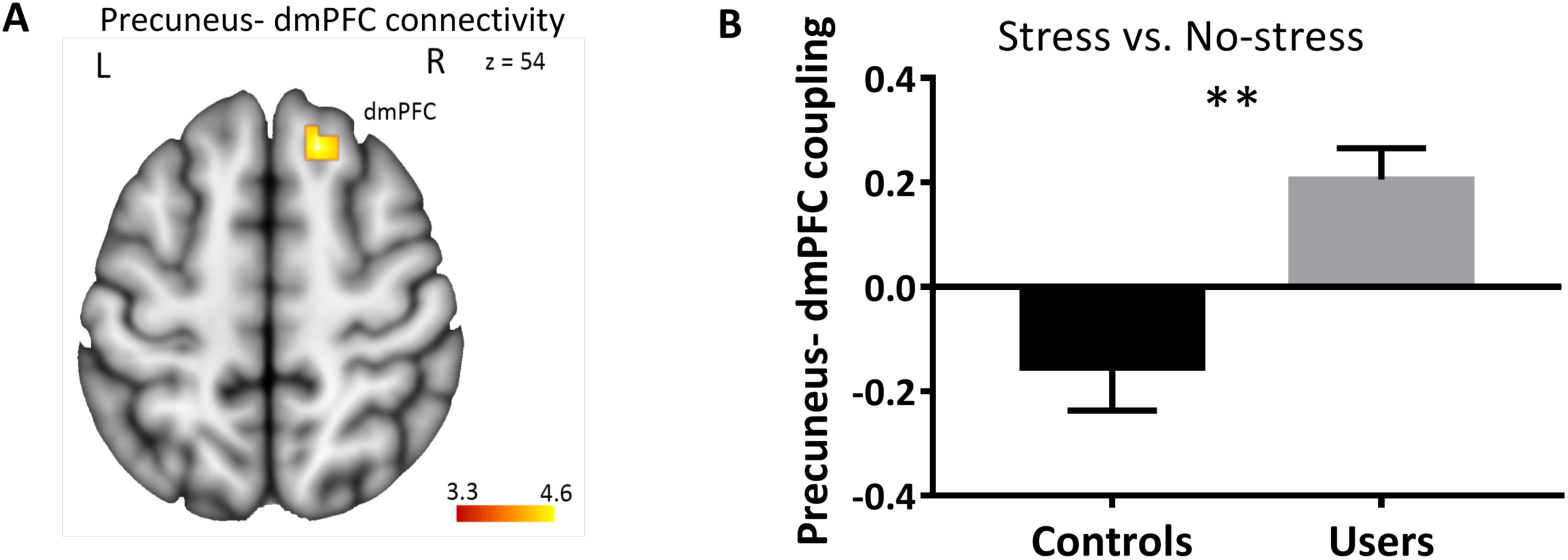
Cannabis users exhibited increased functional connectivity between the precuneus (seed region) and the dorsal medial prefrontal cortex (dmPFC) during stress. **(A)** Location of the precuneus-dmPFC pathway that exhibited group differences, **(B)** extracted connectivity estimates from the pathway for the contrast (stress > no-stress). Image is thresholded at *p* < 0.001, uncorrected. Error bars reflect the SEM. \*\**p* < 0.01. Laterality [left (L), right (R)].

## Discussion

The present study examined psycho-social stress processing in cannabis-dependent males using an adaptive mental arithmetic task accompanied by negative social evaluation. During psycho-social stress exposure, but not the no-stress condition, cannabis users demonstrated decreased performance relative to controls, despite normal stress experience and cardiovascular stress reactivity. On the neural level stress-related performance deteriorations in cannabis users were accompanied by decreased precuneus activity and increased connectivity of this regions with the dmPFC.

In line with previous studies, the experimental task successfully induced stress in both groups as indicated by increased subjective stress experience and cardiovascular activity.^43^ While no differences in cardiovascular and subjective stress reactivity were observed, cannabis-dependent users demonstrated significantly lower arithmetic task performance during stress-induction. Previous studies reported altered stress reactivity in alcohol^44^ and nicotine users^45^ and elevated levels of anxiety and depression.^46^ Cannabis users and controls in our sample were comparable regarding these potential confounders, suggesting specific effects of cannabis dependence on stress reactivity. Both groups exhibited high and comparable performance in the absence of stress indicating comparable baseline cognitive performance. In line with previous studies,^47^ stress increased cannabis craving in dependent subjects, confirming the important role of stress as a driving factor of dependence and relapse.^15^ Against our expectations, however, the groups did not differ with respect to subjective stress experience (however, one study reporting a normal distress experience in cannabis users during social exclusion),^48^ cardiovascular indices as well as self-perceived performance or task enjoyment. Together this suggests that while stress induction and the perception thereof may be intact in dependent cannabis users, psycho-social stress increases cannabis craving and leads to marked deteriorations in cognitive performance.

On the neural level, lower performance in the cannabis group was accompanied by an attenuated stress-related increase in precuneus activity. The precuneus, located in the posteromedial parietal lobe, is considered to play a central role in a range of highly integrated tasks, including basic cognitive (i.e. mental arithmetical performance) as well as social cognitive, particularly self-referential and mentalizing, processes.^49,50^ The precuneus has received little attention in neurobiological models of addiction. However, an increasing number of studies reported altered precuneus activation in chronic cannabis users during cue-induced craving ^51^ as well as cognitive processing in emotional and social contexts such as risky decision making,^52^ suppression of emotional distractors,^53^ evaluation of episodic memory episodes, or mentalizing.^54^

Given the stronger engagement of parietal regions, including the precuneus, with increasing difficulty of mental arithmetic operations,^49^ additional recruitment of the precuneus may have attenuated stress-related task deteriorations in the controls. In contrast, decreased task performance during stress in cannabis users may be linked to failure of compensatory precuneus recruitment. Moreover, decreased stress-related precuneus recruitment was accompanied by increased functional connectivity of this regions with the dmPFC. The dmPFC is involved in both, mental arithmetic performance as well as regulation of negative affect.^49,55^ Increased interactions with the dmPFC may thus reflect a deficient compensatory attempt to maintain task performance or a successful compensation of negative emotional experience during performance deterioration, with the lack of between-group differences in emotional distress arguing for the latter.

The observed psycho-social stress-specific performance impairments in cannabis-dependent subjects align with previous studies reporting impaired cognitive performance in cannabis users in the context of negative emotional^20^ and social information.^56^ Together, these findings suggest an association between chronic cannabis use and deficient integration at the intersection between emotional and cognitive processes. The impact of stress on addiction is multifactorial, and deficient stress regulation has been determined as a risk factor for the escalation of cannabis use and dependence.^13–15^ The retrospective nature of the present study does not allow to disentangle predisposing factors from consequences of chronic cannabis exposure or addiction-related maladaptation in stress regulation. Therefore, impaired performance under stress as well as associated neural alterations may alternatively reflect a predisposing deficit for the development of cannabis dependence or changes in stress-related cannabinoid signaling due to cannabis exposure-related adaptations. From a clinical perspective, the present findings emphasize the detrimental impact of psycho-social stress on craving and cognitive performance in cannabis dependent individuals. As such, exposure to psycho-social stress may promote relapse, impair cognitive performance and ultimately interfere with social and occupational rehabilitation. Therapeutic approaches that aim at improving coping strategies and increase stress resilience in cannabis dependent individuals may therefore represent a promising strategy.

### Limitations and Conclusion

Findings need to be considering in the context of limitations: (1) Only male subjects were included, and the generalizability to female cannabis-dependent users remains to be determined. (2) The groups differed in occasions of ecstasy use. Although a previous study indicates that emotional dysregulations in low-dose ecstasy users are predicted by cannabis rather that ecstasy use,^57^ we cannot rule out a potential contribution on the observed effects. (3) Groups were matched with respect to nicotine use, however the contribution of complex interactive effects between nicotine and cannabis cannot be ruled out. Functional connectivity differences of the precuneus were determined using an exploratory analysis and need to be interpreted with caution.

Overall, the present study provides first evidence for stress-induced cognitive performance deficits in cannabis users. Importantly, deficits were observed specifically for acute stress in contrast to normal performance under no-stress and normal perceived stress intensity. The neural data suggest that deficient stress-related dynamics of the precuneus may mediate these effects.

## Supporting information

supplemental materials and results

