## supplemental materials and results for "Impaired cognitive performance under psycho-social stress in cannabis-dependent males is mediated by attenuated precuneus activity"

Zhao et al.,

### Supplemental Materials

### Data quality assessment and exclusion of participants

Two cannabis users were excluded due to excessive use of other illicit substances. After initial data quality assessments, data from four cannabis users and three controls were excluded due to excessive head movement (> 3.5 mm) during fMRI. Two controls were excluded because they consistently rated subjective stress during the control as well as stress condition very low (stress experience was assessed using a 1-8 scale; one subject consistently rated the stress experience as 1 the other consistently as 2, indicating that the paradigm did not induce stress in these subjects). The final dataset thus included n = 28 dependent cannabis users and n = 23 healthy controls. Details are additionally visualized in the **Figure S1**.

#### MRI data acquisition

Data were acquired on a Siemens Trio 3T MRI system (Siemens, Erlangen, Germany). Functional data was acquired using a T2* echo-planar imaging (EPI) BOLD sequence [repetition time (TR) = 2500 ms, echo time (TE) = 30 ms, 37 slices, slice thickness = 3.0 mm, no gap, voxel size = 2 × 2 × 3 mm, flip angle = 90°, field of view = 192 mm]. To exclude subjects with apparent brain pathologies and facilitate normalization high-resolution T1-weighted structural images were acquired (TR = 1660 ms, TE = 2540 ms, 208 slices, field of view = 256 mm, voxel size = 0.8 × 0.8 × 0.8 mm).

#### fMRI data processing

fMRI data were analyzed using SPM12 (Wellcome Department, London, UK, http://www.fil.ion.ucl.ac. uk/spm/software/spm12). The first five volumes were discarded to achieve magnet-steady images. Next, slice time correction was employed to correct for slice acquisition time and images were realigned using a six-parameter rigid body algorithm to correct for head movement. Subsequently, images were normalized using a two-step procedure that included co-registration with the T1 image, segmentation of the T1 image and application of the resulting transformation matrix to the functional time-series. Images were written out at 3 × 3 × 3 mm resolution and smoothed with a Gaussian kernel (FWHM, 8 mm).

#### fMRI BOLD level Results

In line with previous studies, the paradigm induced widespread activity in psycho-social stress networks encompassing middle frontal regions, precuneus and posterior cingulate cortex (family-wise error correction $p < 0.05$) (see also Dedovic et al., 2009; Wang et al., 2005). See also **Figure S2** showing the overlap between the stress network in the present study and the ROI mask generated from the Eckstein et al., 2014 sample.

**Figure S1** Flow Charts for participants.

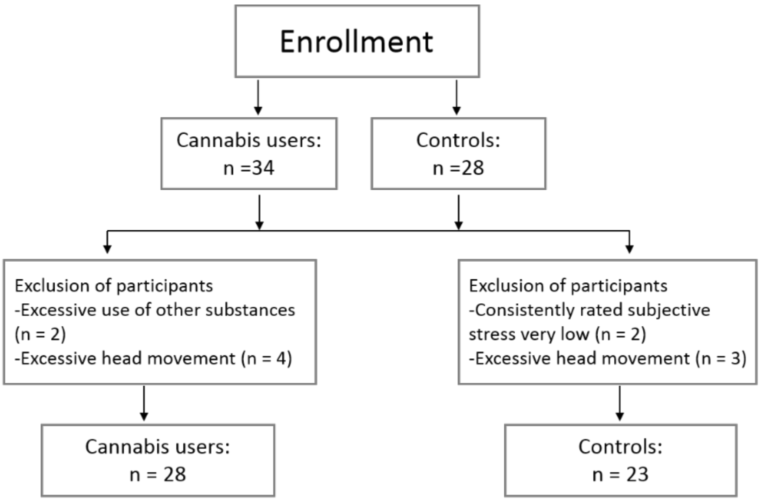

**Figure S2** Stress-related network in the current sample.

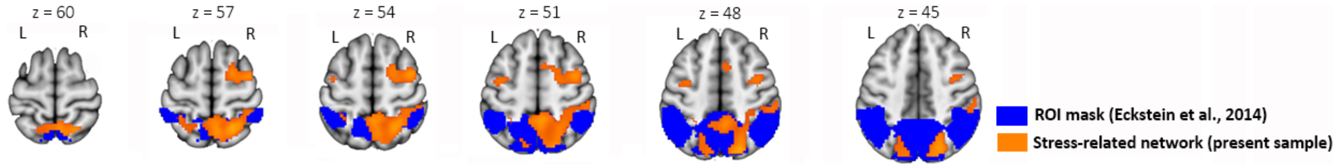
